# Diagnosing Bacterial Vaginosis with a Novel, Clinically-Actionable Molecular Diagnostic Tool

**DOI:** 10.1101/334177

**Authors:** Joseph P. Jarvis, Doug Rains, Steven J. Kradel, James Elliott, Evan E. Diamond, Erik Avaniss-Aghajani, Farid Yasharpour, Jeffrey A. Shaman

**Author notes:** Address correspondence to Jeffrey A. Shaman,.

## Abstract

Bacterial vaginosis is a common condition among women of reproductive age and is associated with potentially serious side-effects, including an increased risk of preterm birth. Recent advancements in microbiome sequencing technologies have produced novel insights into the complicated mechanisms underlying bacterial vaginosis and have given rise to new methods of diagnosis. Here we report on the validation of a quantitative, molecular diagnostic algorithm based on the relative abundances of ten potentially pathogenic bacteria and four commensal Lactobacillus species in research subjects (n = 172) classified as symptomatic (n = 149) or asymptomatic (n = 23). We observe a clear and reinforcing pattern among patients diagnosed by the algorithm that is consistent with the current understanding of biological dynamics and dysregulation of the vaginal microbiome during infection. Using this enhanced assessment of the underlying biology of infection, we demonstrate improved diagnostic sensitivity (93%) and specificity (90%) relative to current diagnostic tools. Our algorithm also appears to provide enhanced diagnostic capabilities in ambiguous classes of patients for whom diagnosis and medical decision-making is complicated, including asymptomatic patients and those deemed “intermediate” by Nugent scoring. Ultimately, we establish CLS2.0q as a quantitative, sensitive, specific, accurate, robust, and flexible algorithm for the clinical diagnosis of bacterial vaginosis – importantly, one that is also ideal for the differential diagnosis of non-BV infections with clinically similar presentations.

## Introduction

Bacterial vaginosis (BV) is widely considered to be the most common vaginal disorder among women of reproductive age (1). In a sample of self-collected vaginal swabs from 3,700 women, the prevalence of BV was estimated at 29% in the general population of women aged 14-49 years and 50% in African-American women (2). Typically, affected patients present with inflammation of the vagina that can result in discharge, itching, and pain. That said, clinical presentation can vary considerably, with asymptomatic infections being noted with great regularity (2). A wide variety of risk factors for BV have been noted, including sexual activity (3,4), the presence of other sexually transmitted infections (5), as well as race and ethnicity (2). Important complications of BV infection include an increased risk of preterm delivery in pregnant women (6) and of HIV acquisition and transmission (7). In some women, clearance of the infection, even with standard of care, is difficult to achieve, often resulting in chronic or recurrent symptoms (8).

Improved strategies for detecting BV infection have been the subject of much discussion, due to the reduction in the quality of life that can result from an active infection and the reasonably high prevalence of BV in women (9). Recent advances in technology have driven interest in DNA-based, quantitative, diagnostic algorithms that are theoretically quicker to perform than culture, more accurate in their results, and are potential candidates as point-of-care devices. However, the seemingly complex biology and dynamics of BV infection, along with important limitations in “gold-standard” approaches, have complicated these efforts.

One confounding factor is the consistent observation that BV is associated not with a single causative pathogen, but rather with a broad disruptive shift in the microbial population: from a healthy *Lactobacilli*-dominated environment to one largely overtaken by organisms such as *Gardnerella*, various mycoplasma, and a heterogeneous collection of facultative anaerobes (10). With this shift come changes in the underlying interactions among different organisms, changes in the metabolites produced by the local bacterial community, and changes to the host’s immune response. Together, these factors produce key signs and symptoms of infection. This dynamic mechanism of disease manifestation results in a condition with all the standard hallmarks of a complex etiology. For example, BV presents clinically in numerous ways, and individual symptoms can be the result of multiple systematic shifts in the vaginal microbiome in favor of a variety of pathogenic organisms. Sometimes women experience no clinical symptoms; in fact, a significant percentage of affected patients are asymptomatic (11,33). This statistic is particularly important in the context of pregnancy, where it has been shown that BV infection, whether symptomatic or not, is associated with preterm birth (6).

In addition, the vaginal microbiota has recently been shown to be longitudinally dynamic in healthy individuals; that is, changes are seen to occur in near-real time over days and weeks (12). The impact of microbial population flux on the maintenance of bacterial community homeostasis, on the resolution of symptoms post-infection, and on the repression of symptoms in asymptomatic women is not well understood. Nevertheless, an oft-changing microbiota poses difficulties for accurate diagnosis, since efficient molecular testing depends on samples taken at a singular point in time.

Given the complexity of the dynamics that lead to clinical presentation, it is unsurprising that diagnosis of BV infections has historically been a challenge. While Gram staining plus Nugent scoring (13) is considered the gold-standard, it is a technique fraught with problems: 1. Nugent requires expertise at staining and scoring morphotypes; 2. several clinically significant microbes, either of variable morphology or lacking a cell wall, are regularly misidentified or overlooked (9); and 3. some 20% of symptomatic subjects score between 4 and 6, rendering their status “intermediate”—an ambiguous diagnosis that is not helpful for the healthcare provider (14) and may lead to inappropriate clinical intervention, or, conversely, to no intervention when it is actually needed. Given the complexity of microbiome alterations now known to be associated with disease, the second of these shortcomings may account for Nugent’s reportedly poor sensitivity and specificity (13). Finally, Nugent scoring has also been shown to occasionally miss differential diagnoses such as *Trichomonas vaginalis* infections (15) with potentially problematic effects on patient well-being due to inappropriate therapy.

An older and more commonly used diagnostic tool is the application of Amsel’s criteria. This test, first described in 1983, requires the satisfaction of at least three of four criteria to qualify for a BV diagnosis: elevated pH (>4.5), characteristic discharge, fishy odor upon the addition of 10% KOH to a slide prep, and the microscopic visualization of “clue cells” (smaller bacteria adhering to epithelial cells) in a wet mount (16). However, while easy to implement in theory, Amsel’s criteria are reportedly underutilized in clinical settings (17) and are technically limited by the lack of consideration of the abundance of healthy *Lactobacillus* species (18).

These and other acknowledged failings of the gold-standard diagnostic techniques for BV have recently fueled a search for alternatives leveraging new technology. Today, there are a variety of alternatives in the marketplace that claim ease of testing, enhanced sensitivity and specificity, as well as reduced cost. Relatively few of these can be used at the point-of care (PoC) (19,20); rather, most available tests require sending a sample – often in the form of a vaginal swab – to a laboratory for testing. Though not as expedient as a PoC diagnostic, at 1 to 2 days for a result after sample transport and testing, versus minutes to hours for a PoC result, the referencing of samples to an external laboratory offers several key advantages. Most notably, some clinical laboratories can simultaneously screen samples for a broad range of microbial targets covering multiple conditions, not merely BV. The ability to offer a differential diagnosis is a key advantage for symptomatic patients, since infections other than BV can present with very similar signs and symptoms. Many labs can also generate bacteria-specific presence/absence or semi-quantitative information, while also detecting difficult-to-culture, anaerobic, and commensal microorganisms. Such data offer the potential for healthcare providers, who are often faced with interpreting ambiguous signs and symptoms, to receive greater diagnostic clarity. Similarly, they can be alerted to mixed/co-infections, a situation reported to affect 20% of BV+ patients (21). Finally, providers can potentially receive specific recommendations for therapy (e.g. antibiotic regimens or combinations of treatment options specific to the pathogen(s) detected). While in no way a replacement for a thorough physical examination and detailed capture of patient history (22), such molecular-based screening holds great promise for the enhancement of the current standard-of-care.

Here we present the refinement and performance evaluation of an existing quantitative molecular diagnostic for BV that eliminates the challenges demonstrated by Amsel and Nugent tests, while also providing an accurate diagnosis via genus and species-specific information for both pathogenic and non-pathogenic bacteria. The test, based on real-time PCR, is offered in a highly customizable format, allowing individual laboratories to add or subtract content as future needs and research developments dictate. The approach also permits simultaneous screening of an assortment of other vaginal pathogens with no additional labor or consumable requirements, thereby enabling differential diagnoses – an added value of this test over current PoC solutions.

## Materials and Methods

### Diagnostic Panel Design

As part of previous BV diagnostic algorithm development efforts, we conducted a comprehensive review of published studies on normal and pathogenic vaginal microbiota. Our survey included literature characterizing the microbiomes of BV+ and BV-women via next-generation sequencing (12,17,23,24), a survey of molecular-based diagnostic tool development for BV (1,9,14,25,26,27), and expert reviews of both of these subjects (28,29,30,31,32). We found that numerous studies have shown that the amounts of particular bacteria – and not merely their presence or absence – play a role in the development and/or persistence of BV (1,9,14,17,27,28,31,34,35). Ultimately, fourteen BV-associated microorganisms (TABLE 1) that would theoretically maximize diagnostic specificity while remaining cost-effective to the clinical laboratory were selected for inclusion in the new diagnostic test. TaqMan®-based assays were designed with the assistance of ThermoFisher’s bioinformatic pipeline to detect bacterial-specific genes down to the species level, reported to be present at one copy per microbe, a fact important for the relative abundance calculation of each microbial target. The assays were subsequently spotted into the through-holes of a custom-designed TaqMan OpenArray plate for vaginal microbiota investigations (available from ThermoFisher Scientific).

**TABLE 1:** Microorganisms included in the CLS2.0q molecular assay for Bacterial Vaginosis diagnosis. After extensive literature review, these ten likely-pathogenic and four commensal bacteria were chosen as targets associated with bacterial vaginosis. The relative amounts of each of these bacteria present in a patient sample inform the CLS2.0q diagnostic.

| Pathogenic | Commensal lactobacilli |
| --- | --- |
| *Atopobium vaginae* | *Lactobacillus crispatus* |
| *BVAB2\** | *Lactobacillus gasseri* |
| *Gardnerella vaginalis* | *Lactobacillus iners* |
| *Megasphaera 1* | *Lactobacillus jensenii* |
| *Megasphaera 2* |  |
| *Mobiluncus curtisii* |  |
| *Mobiluncus mulieris* |  |
| *Mycoplasma hominis* |  |
| *Prevotella bivia* |  |
| *Ureaplasma urealyticum* |  |

\*Bacterial Vaginosis–Associated Bacterium type 2

Since bacterial vaginosis, candidiasis, and trichomoniasis are all indicated in vaginitis, present clinically with similar signs and symptoms, and are often observed in co-infected patients, assays specific to these infections were added to the panel as an aid to differential diagnoses. Furthermore, because *Candida albicans* infections are treated differently than non-*albicans* infections, five specific yeast species were also included in the final panel design.

### Algorithm Development and Study Population

During the original algorithm design phase, several statistical and algorithmic approaches were applied to a sample of patients (410 antibiotic-naïve, including 25 asymptomatic women showing no clue cells on analysis for Amsel’s criteria) and evaluated with respect to sensitivity and specificity. These included a number of multivariate linear and mixed models as well as a variety of other techniques. In general, linear regression techniques performed poorly with respect to sensitivity (i.e., the ability to identify BV+ individuals), but performed fairly well with respect to specificity (i.e., the ability to exclude BV as a diagnosis). Alternative strategies, such as those that produced the final CLS1 algorithm, showed improved overall performance (especially increased sensitivity), albeit with slightly reduced specificity. Ultimately, the highest performing quantitative model using measures of relative abundance was chosen for validation, as opposed to one based solely on presence/absence data.

Subsequently, a total of 172 research subjects (149 symptomatic and 23 asymptomatic women) were recruited into a clinical study aimed at validating the existing, proprietary diagnostic algorithm (CLS2.0q). All were enrolled at the Maternity & Infertility Institute (San Fernando, CA). Patients younger than 18 years old or those who had taken antibiotics in the previous 30 days were excluded. Patient ethnicity, age, pregnancy status, menstruation status, history of recurrent bacterial vaginosis and candidiasis, and method of birth control were recorded. Nugent scoring was also performed on each. The average age of enrollees was 30.4 years; 91.8% self-classified as Hispanic/Latina.

### Sample collection and analysis

As in the population used for algorithm development, three samples from the lateral vaginal wall and posterior fornix or a blind vaginal swab were collected in the validation population using COPAN ESwab™ with flocked nylon fiber tips (36). Two of the swabs were transferred into 1 mL of modified Liquid Amies Transport medium and shipped at room temperature to the laboratories for DNA processing. One swab was used to transfer collected sample material onto each of three slides.

One slide was examined for *Trichomonas vaginalis* and clue cells using standard microscopy techniques. The second slide was examined using the whiff test by adding a drop of 10% KOH solution and checking for a characteristically strong fishy odor. The third slide was sent to Primex Clinical Laboratories (Van Nuys, CA) for independent evaluation by a pathologist and technician blinded to the clinical presentation of the sample’s donor.

The pathologist and technician applied Nugent scoring methods (13) to a smear of a sample collected from each patient. Briefly, the swab was rolled on a glass slide; the smear was then heat-fixed, and Gram stained. The smear was subsequently evaluated for the following morphotypes (1000x magnification under oil immersion): large Gram-positive rods (*Lactobacillus* morphotypes), small Gram-variable rods (*G. vaginalis* morphotypes), small Gram-negative rods (*Bacteroides* species morphotypes), and curved Gram-variable rods (*Mobiluncus* species morphotypes). Nugent scoring was used to diagnose bacterial vaginosis. For samples with a 4-6 “intermediate” score, the presence of three of four the Amsel’s criteria (grayish-white homogeneous vaginal discharge, pH≥4.5, positive whiff test, and presence of clue cells) was used as the criterion to diagnose BV.

For the molecular assays, DNA was isolated using MagMAX™ Express 96 semi-automated sample prep system in combination with the MagMAX™ DNA Multi-Sample Ultra Kit (ThermoFisher Scientific, Waltham, MA) and standard protocols. Next, real-time PCR was performed on multiple replicates using 2.5 uL of 2X OpenArray® Gene Expression Master Mix combined with 2.5 uL eluted DNA in a 384-well plate and added to the OpenArray plate using the AccuFill™ (ThermoFisher Sc.) automated pipetting instrument. OpenArray plates were cycled on the QuantStudio 12K Flex OpenArray Real-Time PCR System (ThermoFisher Sc.) according to the manufacturer’s recommended conditions. Following cycling, data were analyzed and exported using QuantStudio software. Samples were only counted as true positives when at least 50% of technical replicates for a given microbe-specific Assay yielded a Crt value ≤31. Additionally, any sample in which the 16S rDNA control failed to achieve a minimal threshold of 21 was rejected, since higher values suggest inadequate sample collection. Relative quantities for each BV-associated bacterial assay were calculated based on Crt values for each of the targets.

## Results

Clinical samples from 149 symptomatic and 23 asymptomatic women were tested for the relative abundance of the 14 distinct bacterial species (TABLE 1) that were chosen to provide a clinically-significant representation of the vaginal microbiome. Based on these data, patients were diagnosed as either BV+ or BV-using a previously developed, proprietary, quantitative molecular diagnostic algorithm, CLS2.0q (FIG 1a). Sensitivity and specificity were assessed with respect to the gold-standard Nugent test and found to be 93% and 90%, respectively.

**FIG1:**
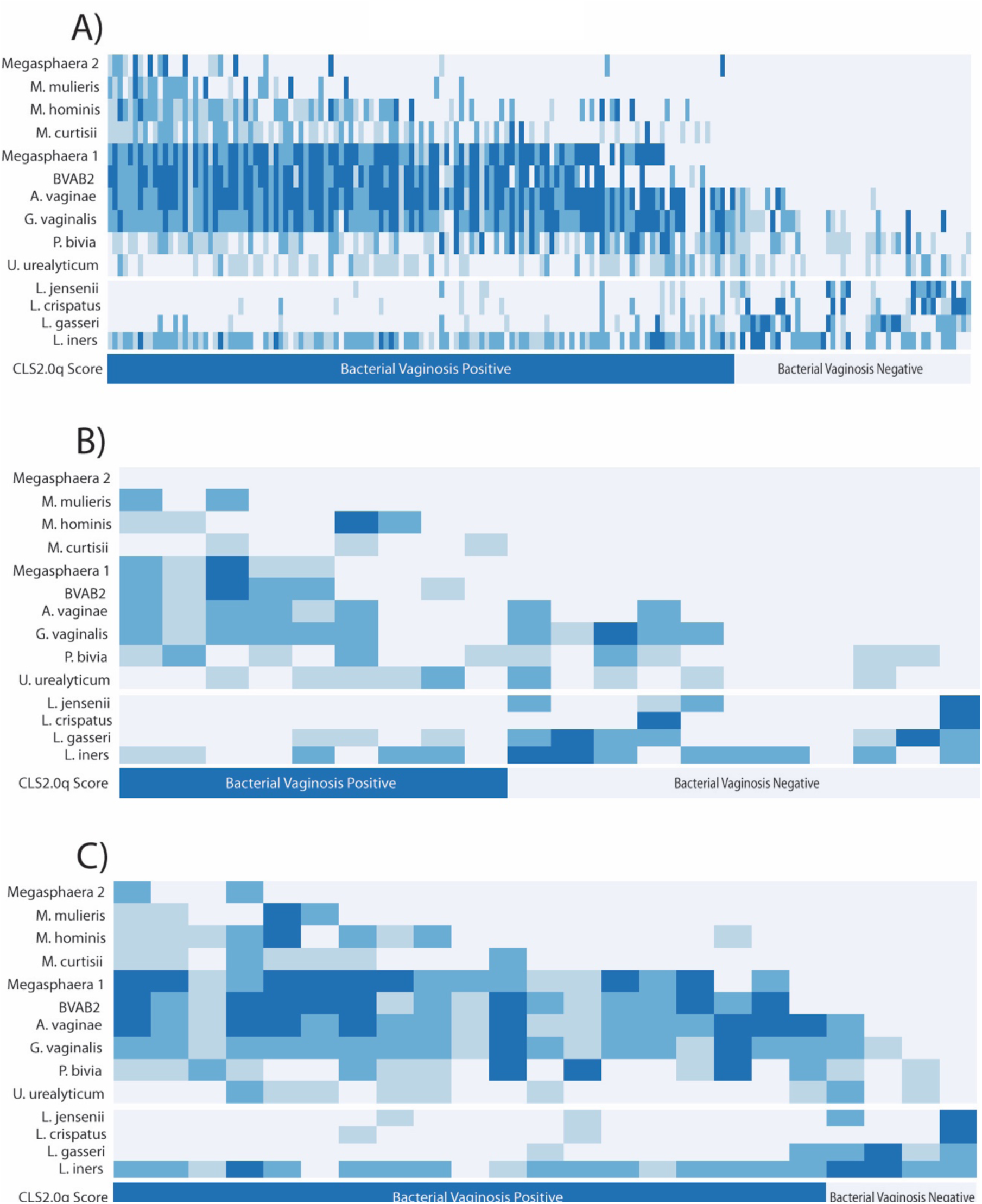
Relative bacterial abundance in the validation data set (N = 172), and its role in CLS2.0q diagnosis of bacterial vaginosis in vaginal microbiome. (A) Relative bacterial abundance (rows) of pathogenic (top section), commensal Lactobacilli (middle section) in women (columns) diagnosed as BV+ and BV-(bottom section) by CLS2.0q in the validation dataset. No single organism perfectly differentiates BV+ and BV-patients. Rather, BV+ subjects show relatively high levels of multiple pathogenic bacteria especially Megasphaera 1, BVAB2, *A. vaginae*, and *G. vaginalis* (top-left) and relatively low levels of commensal Lactobacillus species (middle-left). In contrast, BV-subjects show relatively high abundances of multiple Lactobacillus spp. (middle-right) and a concurrent lack of pathogenic bacteria (top-right), especially Megasphaera1 and BVAB2. Together, these mirror-image patterns constitute the primary signal detected by CLS2.0q in diagnosing BV in clinically symptomatic and asymptomatic women. (B) Relative bacterial abundance and CLS2.0q diagnosis of BV for samples with a Nugent categorization of “Intermediate” (N = 20). A strikingly similar pattern as described above is associated with BV diagnosis by CLS2.0q. (C) Relative bacterial abundance and CLS2.0q diagnosis of BV for asymptomatic individuals (N = 23). A similar and consistent pattern as described above is also associated with BV diagnosis by CLS2.0q.

Several clear patterns emerged when comparing diagnostic outcomes with observed variation in bacterial abundance. First, BV+ women were found to have relatively high levels of multiple pathogenic organisms. No single organism perfectly differentiated between BV+ and BV-patients; rather, the combined presence of several microbes (especially Megasphaera 1, BVAB2, *A. vaginae*, and *G. vaginalis*) contribute to a clear signal of disease. Furthermore, a concurrent paucity of non-pathogenic, commensal *Lactobacillus* species in BV+ patients was also observed. Together, these patterns reinforce one another in the diagnosis of BV by CLS2.0q consistent with *a priori* biological expectations and previous analyses (FIG 1a). In contrast, women diagnosed as BV-show precisely the opposite pattern. Specifically, BV-patients generally present with relatively high abundances of multiple putatively-beneficial lactobacillus spp. and a concurrent lack of pathogenic bacteria, especially those most often found at high levels in BV+ patients (e.g. Megasphaera1, BVAB2; FIG 1a). Together, these patterns constitute the primary signal detected by CLS2.0q in diagnosing BV in clinically symptomatic and asymptomatic women.

Interestingly, the non-pathogenic bacterium *L. iners* is relatively abundant in many patients regardless of their diagnosis. This suggests either that the presence and abundance of *L. iners* is not relevant to BV diagnosis or, more interestingly, that it plays differing roles in the disease-versus healthy-state communities of bacteria. Indeed, previous studies have suggested that *L. iners* acts as a “sentinel” or “transition” species early in the course of pathogenic infection (37) and may persist at relatively high abundance thereafter. Regardless, this pattern of abundance across patients likely does not contribute significantly to the diagnostic signal. Similarly, *G. vaginalis, P. bivia*, and *U urealyticum* are often found in both BV+ and BV-patients. This also suggests they do not contribute strongly to the diagnosis of disease. Rather, this pattern likely reflects the natural dynamics of a community of bacteria at equilibrium with beneficial types, keeping the explosive growth of pathogenic organisms in check.

Several, relatively rarely observed, pathogenic bacteria (e.g. Megasphaera 2 and *M. mulieris*) were included in testing. Despite this rarity, it is clear that they contribute to the overall patterns discussed above. In point of fact, their strongest contribution to diagnosis is likely their conspicuous absence in BV-patients.

Over the course of data analysis, 20 subjects with intermediate Nugent scores were identified. Such patients represent an ambiguous class in current, diagnostic practice with unclear guidelines for treatment. Interestingly, via the mutually reinforcing patterns described above, CLS2.0q appears to provide clear resolution for such patients (Fig. 1b). Specifically, roughly half (n=9) are diagnosed as BV+ by the algorithm and half (n=11) are diagnosed as BV-.

Upon further inspection, all asymptomatic women in the validation dataset (N = 23) show some evidence of BV by Nugent analysis. Specifically, 16 score as “High” and the remaining seven as intermediate upon scoring. None score as “Low.” Interestingly, four of these women are diagnosed as BV+ by CLS2.0q, all of which are Nugent intermediate but were designated by the microscopist as having “few” clue cells.

Assays for candidiasis, trichomoniasis, and five specific *Candida* species were included in the final panel design to enable differential diagnosis for these infections due to their clinically similar presentations (TABLE 2).

**TABLE 2:**
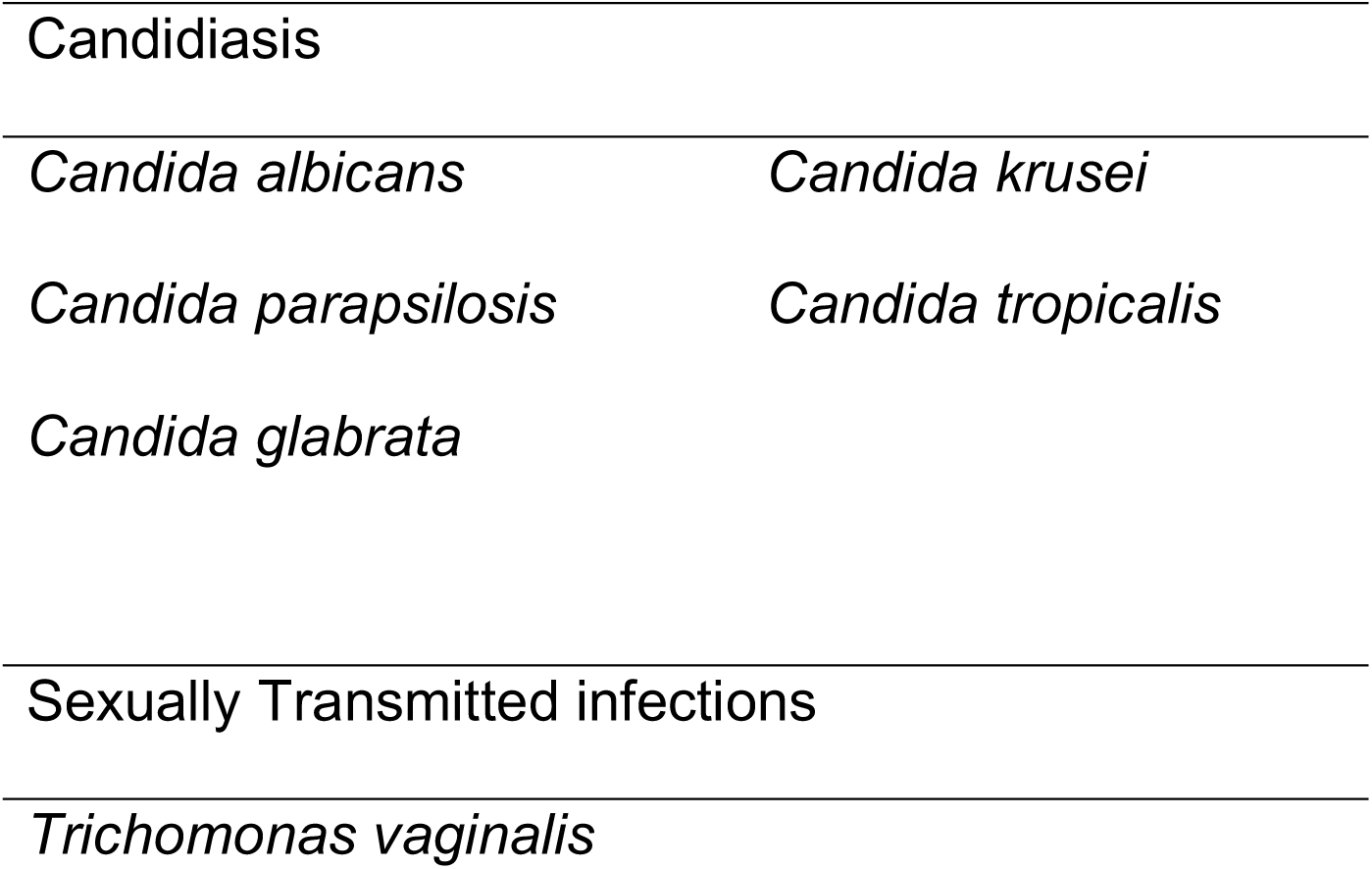
Additional organisms included in the CLS2.0q molecular assay for differential diagnosis. After extensive literature review these organisms were chosen as targets associated with infections which present with clinically similar signs and symptoms to BV. The presence of any of these informs the process of differential diagnosis.

## Discussion

With a measured prevalence between 29% and 50% in women of reproductive age, there are a significant number of women who are negatively impacted by bacterial vaginosis infections each year. Infection is associated with serious potential side effects including increased risk of preterm birth in women affected during pregnancy. However, the complex etiology of the condition — the way multiple distinct combinations of bacteria are associated with a disrupted vaginal microbiome — has only recently begun to be appreciated. Furthermore, patients present with a wide range of clinical symptoms from entirely asymptomatic to showing multiple signs and symptoms. Others are actually infected with other pathogens that cause similar signs and symptoms. The dynamic and highly interactive nature of the underlying cause of disease and complicated clinical presentation makes accurate diagnosis—and differential diagnosis—a challenge.

As such, improved strategies for detecting BV infection have been a subject of much discussion. Recent advances in sequencing technologies, the very same that have begun revealing the biology responsible for disease, also show great promise for improving diagnosis. They even open the door to the development of personalized treatment plans based on the actual combination of bacteria at both increased and decreased relative abundance.

Here we describe a novel, molecular diagnostic based on quantitative measurement of the relative abundance of multiple pathogenic and commensal organisms of the vaginal microbiome. It does not rely on culture, microscopy, or subjective analysis of patterns of staining. Rather, mutually reinforcing patterns in the observed levels of each of the assayed microorganisms accurately capture the biological complexities of the vaginal microbiome and clearly differentiate BV positive and BV negative patients.

The complicated biology underlying BV infections suggests that accurate methods for diagnosis must take into account the balance of multiple organisms simultaneously. Historically, methods such as Amsel’s criteria have done this via numerous clinical observations and tests, or by bacterial staining and other laboratory-based techniques in the case of Nugent scoring.

However, these methods utilize presence/absence determinations for some diagnostic features and for others provide merely a semi-quantitative categorization of patients. These limitations result in poor resolution by the gold-standard approach. Specifically, Nugent scoring provides a somewhat-ambiguous finding of “Low,” “Intermediate,” or “High” rather than a more concrete diagnosis of “BV+” or “BV-.”

The diagnostic method we validate here appears to much more accurately capture the complex biology of BV compared to existing alternatives. It identifies a clear reinforcing pattern of abundance (see Fig. 1a) of multiple pathogenic and commensal *Lactobacilli* species that provides a clear and accurate diagnosis, even in diagnostically ambiguous “Intermediate” patients (Fig. 1b). The method also provides improved sensitivity and specificity over current standard-of-care diagnostic tools. Interestingly, these metrics likely underestimate the number of true positive and true negative individuals, since the algorithm was necessarily compared to a standard with known flaws. It is possible that these flaws produce an upper limit on performance that is unrelated to the clear signal produced by CLS2.0q. The actual sensitivity and specificity of the CLS2.0q diagnostic algorithm may therefore be greater than reported.

In addition to providing clarity for the ambiguous Nugent “Intermediate” subjects, CLS2.0q also appears to perform well in asymptomatic patients. This is especially valuable as it is well documented that a majority of BV cases present with no clinical symptoms at all. Interestingly, most of the asymptomatic patients involved in the validation were diagnosable by CLS2.0q as BV+ (FIG 1c). Again, via clearly interpretable biological pattern recognition, CLS2.0q appears to aid in the accurate diagnosis of such patients as well, though additional studies focused on this subgroup would be necessary to confirm this observation.

In addition to the improved capture of biologically-consistent, patient-specific patterns of dysregulation in the vaginal microbiome, CLS2.0q also incorporates several technical improvements over existing methods. For example, together with the detection technology, it is capable of identifying both rare and difficult-to-culture organisms that are currently missed by standard diagnosis methods. Additionally, OpenArray technology provides several key advantages over previous approaches. These include increased flexibility of microbial target choices, improved accuracy of microbial identification, and high-throughput capability. They also allow for the refinement of platform content as future research continues to reveal the underlying dynamics of infection.

This successful validation of the CLS2.0q algorithm was nonetheless limited in several ways. The first is the somewhat small overall sample size in the validation set of 172 women. However, though the validation dataset may seem limited, the original training set included 410 symptomatic and asymptomatic women, and all findings during the original algorithm development were very similar to those presented here (data not shown). The second limitation is this study’s somewhat biased sample of patients. Whereas the majority of subjects in this dataset self-identify as Latina, the ideal validation would have included representative samples of all ethnicities in order to fully control for known and potential differences amongst groups. An ideal study would also have fully controlled for geography and socio-economic status. However, the availability of subjects and other logistical challenges limited collection to a single location/health system whose population may not be fully representative at the national level.

Another limitation is inherent to the biology of the condition itself, namely the dynamics of the vaginal microbiome in real-time that inevitably introduces variation due to the status of the dysregulation at the time of sampling. While formally unaccounted for in our analyses, the robustness of the algorithm displayed in development and in this validation strongly suggest that we have sampled over a sufficient number of dysregulated states to reflect the majority of the relevant biological variation detectable by the method.

Ultimately, this validation established CLS2.0q as a quantitative, sensitive, specific, accurate, and robust algorithm for the diagnosis of bacterial vaginosis. It is demonstrably superior to gold-standard approaches at identifying the underlying biological dynamics of dysregulation in the vaginal microbiome that underlie the condition. It enables efficient differential diagnosis of multiple other conditions that present in a clinically similar manner. Finally, future work may allow the algorithm to be extended to the data-driven recommendation of personalized treatment regimes, including those that may help circumvent and/or avoid antibiotic resistance. Ongoing research is expected to identify additional organisms of interest for bacterial vaginosis, and CLS2.0q is well positioned to rapidly include this information in future refinements of the algorithm.

It is evident that quantitative methods of diagnosis provide key improvements over current approaches to the diagnosis and treatment of complex infections. It is hoped that the technology employed here will enable point-of-care compatible turnaround times and broaden the impact of these techniques in personalized medicine. In addition to flexibility and efficiency, there is also the potential for these approaches to identify drug-resistant organisms, and thus to suggest specific and personalized treatment regimens for individual patients. Furthermore, these techniques can be used to test for and diagnose other conditions, such as respiratory infections, gastrointestinal dysregulation, and wound healing concerns.

## Acknowledgments

The authors thank ThermoFisher Scientific for their support in obtaining the data used to train and develop the CLS2.0q algorithm.

